# DeepSpaceDB 2.0: an interactive spatial transcriptomics database for large-scale Xenium data exploration

**DOI:** 10.64898/2026.01.15.699623

**Authors:** Vladyslav Honcharuk, Keiko Takemoto, Marvin C. Masalunga, Hongyi Zhao, Diego Diez, Shinpei Kawaoka, Alexis Vandenbon

## Abstract

The 10x Genomics Xenium platform enables high-resolution spatial transcriptomics at single-cell and subcellular scales, but effective reuse of public Xenium datasets is hindered by large data sizes and heterogeneous file formats. We previously developed DeepSpaceDB, a spatial transcriptomics database designed for interactive, in-depth analysis of tissues and tissue microenvironments. Here, we present a major expansion of DeepSpaceDB that integrates large-scale single-cell spatial transcriptomics data generated by the Xenium platform. In this update, we systematically collected 1,539 public Xenium datasets from multiple repositories and processed them through a robust, standardized pipeline that validates, repairs, and harmonizes heterogeneous inputs into a unified representation. To support efficient exploration of these data, we introduced a redesigned DeepSpaceDB interface and complementary Zarr-based storage formats optimized for gene-centric visualization and spatially localized queries, enabling sub-second response times for common interactive operations. The updated platform supports real-time visualization of spatial data and analysis of regions of interest directly in the web browser. Together, this expansion establishes DeepSpaceDB as a unified resource for single-cell spatial transcriptomics, substantially lowering the barrier to accessing, exploring, and reusing large-scale public Xenium datasets.

## INTRODUCTION

High-resolution spatial transcriptomics technologies enable the measurement of gene expression at single-cell and subcellular resolution while preserving spatial context. Among these, the 10x Genomics Xenium platform provides in situ detection of tens of millions to hundreds of millions of transcripts across large tissue sections, generating datasets that combine high molecular resolution with substantial spatial complexity. As public Xenium datasets accumulate, there is an increasing need for tools that support efficient exploration, visualization, and comparison of these data across studies.

Despite the growing availability of Xenium data, their reuse remains challenging. Xenium data is large and complex. Publicly released datasets vary widely in the combination of files provided, with many samples lacking essential data such as the molecule-level transcript tables, cell segmentation outputs, or cell-by-gene count matrices. Even for samples where all data is available, their processing requires a high level of bioinformatics expertise. A number of spatial transcriptomics databases have been published with the aim of making spatial transcriptomics data more easily accessible (Fan, Chen, and Chen 2020, Yuan *et al*. 2023, Xu *et al*. 2024). However, while these databases tend to cover many different platforms, the data exploration functions are very limited. Moreover, they don’t include Xenium samples, presumably because the large numbers of cells pose significant burdens on the visualization functions of these databases. Notable exceptions are the SOAR and HISSTA databases (Li et al. 2025, Yu et al. 2025). SOAR recently added a number of Xenium samples, but its interface features only static low-resolution plots, lacks image data, and is not suitable for interactive exploration. HISSTA contains both single cell and spatial transcriptomics data, including 46 Xenium samples. The samples in HISSTA can be accessed through Vitessce (Keller et al. 2025), allowing visualization of gene expression and clustering results. However, it lacks imaging data, no interactive comparisons between regions can be made, and only processed Seurat objects are available for download and not raw data.

We recently developed DeepSpaceDB, a database that facilitates interactive exploration of spatial transcriptomics data (Honcharuk et al. 2025). For example, DeepSpaceDB allows users to interactively select regions of interest and compare gene expression between multiple selected regions. Whereas DeepSpaceDB originally focused on samples of the Visium platform, we here expand it to include 1,539 samples of the high-resolution Xenium platform and a dedicated Xenium interface that enables fast, interactive exploration of large Xenium datasets. In this manuscript, we describe the design and implementation of the DeepSpaceDB Xenium interface. We outline how public Xenium datasets are collected and harmonized despite heterogeneous file availability, how data are stored using complementary Zarr-based (https://zarr.dev/) representations optimized for gene-centric and spatial queries, and how these design choices enable an intuitive and responsive user interface. Finally, we summarize key properties of the integrated datasets and demonstrate representative use cases for interactive exploration, cross-sample analysis, and database-wide search.

## MATERIAL AND METHODS

### Data collection and quality control

Xenium data was obtained from three public repositories - NCBI GEO, 10x Genomics Website and DBKERO (Barrett et al. 2013, Suzuki et al. 2018). Raw data typically comprises transcriptome data, cell counts, spatial coordinates and fluorescence and/or H&E imaging data. Collected data includes both Xenium v1 samples, which cover gene panels of up to 500 genes, and Xenium Prime 5k samples, which cover approximately 5,000 genes.

The raw Xenium data were highly heterogeneous. Samples from the 10x Genomics Website and DBKERO, and a limited number of GEO samples included all necessary files for directly processing using the SpatialData package or other similar approaches (Marconato et al. 2025). However, the majority of GEO-derived samples had inconsistencies in their data, including different types of compressions, inconsistent file names or different data formats (e.g. csv instead of parquet). In addition, a significant fraction of samples was incomplete, with many samples lacking gene-expression data, spatial coordinate data, or imaging data.

To address these issues, we devised a systematic pre-processing pipeline (Figure 1 and Supplementary section “DeepSpaceDB Xenium Data Processing Pipeline”; https://github.com/vladyslav-honcharuk/deepspacedb-xenium-pipeline). Each sample is first subjected to a validation step that verifies the presence of all required files and checks for format consistency. Missing or malformed files are either recovered (e.g. fixing issues in the index column of the barcodes.tsv file, etc.) or regenerated (e.g. generating cells.parquet files from MEX counts and boundary data) when possible.

**Figure 1.**
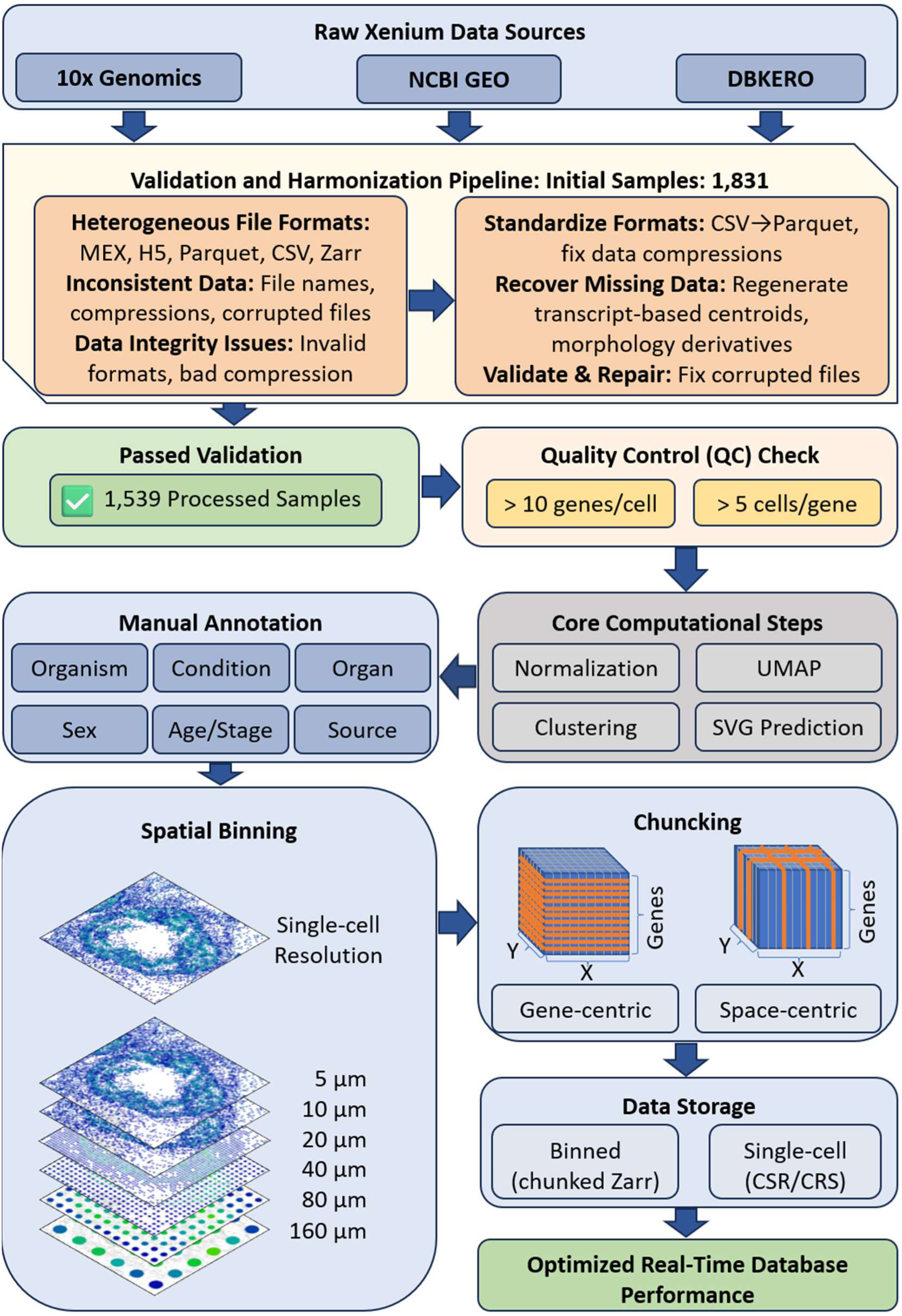
Schematic summary of the data pre-processing pipeline. In brief, raw Xenium data is collected from several sources. Inconsistencies in the available data are corrected as far as possible. Correctly processed samples are normalized, annotated, and processed into gene-centric and space-centric Zarr stores for fast access from the DeepSpaceDB Xenium interface.

Once all files are validated and harmonized, we process them using the SpatialData and SpatialData-io Python packages (versions 0.5.0 and 0.3.0, respectively) (Marconato *et al*. 2025). SpatialData provides a way to standardize data and facilitates downstream analyses such as clustering and quality control. However, because it assumes the standard Xenium raw format with set filenames, it cannot be directly used on samples that deviate from this standard, and requires manual file renaming. Our workflow therefore first attempts to salvage any sample deviating from the canonical formats before it is processed by SpatialData. Of the 1,831 Xenium samples we initially collected, our pipeline was able to successfully process 1,539 samples. These samples were then subjected to an initial quality check using SpatialData and scanpy (Wolf, Angerer, and Theis 2018), including calculating the number of detected genes per cell, and the number of cells in which each gene was detected. Cells with less than 10 detected genes, and genes detected in less than 5 cells were filtered out. A rough quality indicator for each sample was estimated based on database-wide trends (see Supplementary section “Database-wide quality trends”).

In addition, we manually collected annotation data for successfully processed samples, including organism, organ or tissue, medical condition, sex, age or stage, as far as possible. Organ of origin and medical condition annotations were mapped onto the Uberon anatomy ontology (Haendel et al. 2014) and Human Disease Ontology (DO) (Baron et al. 2024).

### Transcriptome data processing

Each Xenium sample was processed through a standardized pipeline implemented in Python using spatialdata, scanpy, and saved as a series of Zarr stores. Following suggested best-practices (Marco Salas *et al*. 2025), data was normalized using scanpy’s normalize_total and log1p functions with default parameters. Dimensionality reduction was performed using Principal Component Analysis (PCA). A k-nearest neighbor graph was constructed on the PCA embedding, which was then used for UMAP dimensionality reduction and clustering using the Leiden algorithm (Traag, Waltman, and Eck van 2019) via scanpy’s neighbors and leiden functions with default parameters. Spatial domains were predicted using BANKSY (Singhal *et al*. 2024) with a Leiden clustering resolution of 0.9, 20 PCA dimensions, λ = 0.8, and 15 spatial neighbors. Spatially variable genes (SVGs) were identified using singleCellHaystack (Vandenbon and Diez 2020, 2023) with default parameters.

Xenium raw data files average 9 GB per sample (range: 1–20 GB), posing a significant bottleneck for real-time data retrieval in an interactive database interface. To enable rapid data access for the database interface, each sample underwent several optimization steps:

#### Spatial binning and chunking

Expression data was prepared at seven spatial resolutions stored as Zarr archives. At single-cell resolution, each cell was represented by its centroid coordinates. Additionally, data was spatially binned at six resolutions (5, 10, 20, 40, 80, and 160 μm bin sizes), with transcript counts summed within each bin. For each resolution, two complementary chunking strategies were implemented: (1) gene-chunked stores (chunks: 1 gene × full spatial extent) optimized for visualizing individual gene expression patterns, and (2) spatial-chunked stores (chunks: all genes × 100×100 spatial bins) optimized for calculating expression across all genes within user-defined regions of interest.

#### Single-cell expression formats

Full single-cell expression matrices were stored in three sparse matrix formats: CSR (cell-wise access), CSC (gene-wise access), and per-gene chunked arrays - to optimize different query patterns.

#### Quality metrics

Comprehensive quality control metrics were calculated on raw count data, including per-cell and per-gene statistics. Multiple data layers (raw counts, normalized, log-transformed) were preserved, and SVG rankings with associated statistics (Kullback-Leibler divergence, p-values as calculated by singleCellHaystack) were saved for each sample.

All processed data were packaged as compressed Zarr stores to enable efficient random access and partial reading required for interactive database queries.

### Cell type predictions

We predicted cell types in all Xenium samples by comparing cluster-level gene expression with reference transcriptomic datasets containing annotated cell types. Briefly, we collected human and mouse single-cell RNA-seq datasets with cell type annotations from the CellxGene database (Megill *et al*. 2021). The data were normalized and converted into pseudobulk reference profiles by averaging gene expression across cells of the same cell type. In addition, each reference cell type was assigned to a smaller set of “broad” cell types. This resulted in a human reference comprising 222 cell types grouped into 18 broad cell types, and a mouse reference comprising 87 cell types grouped into 11 broad cell types. For each Xenium sample, we calculated the Spearman correlation between the gene expression profile of each Xenium cluster and each reference cell type. Each cluster was assigned the cell type with the highest correlation, together with its corresponding broad cell type. Because Xenium datasets are often based on relatively small gene panels and can exhibit substantial technical noise, anatomically implausible predictions were occasionally obtained (e.g., hepatocytes in brain tissue). We therefore applied a simple anatomy-based filter to remove biologically implausible assignments. In DeepSpaceDB, users can inspect both detailed and broad cell type annotations. More details are provided in the Supplementary Methods.

### Spatial neighborhood enrichment analysis

Cell-cell neighborhood enrichment was computed was using squidpy (version 1.6.2) (Palla *et al*. 2022). In brief, in regions of interest selected by the user, first a spatial neighbor graph was built between the cell types with squidpy.gr.spatial_neighbors using k-nearest-neighbor graph (coord_type=‘generic‘, default n_neighs=6). Cell types represented by fewer than 10 cells were excluded prior to the enrichment test. Neighborhood enrichment z-scores were then computed with squidpy.gr.nhood_enrichment using default parameters, with a fixed random seed of 42. Positive z-scores indicate spatial co-localization between the cell type pairs, while negative z-scores indicate spatial avoidance.

### Gene co-expression analysis

In regions of interest selected by the user, the Pearson correlation between a query gene of interest and all genes included in the sample is calculated. Top 10 genes with the highest positive, and the highest negative correlation are visualized in the interface. These genes are also clustered by their mutual Pearson correlation and plotted as a heatmap. Pearson correlation values can also be downloaded in .csv format.

### Image data processing

For samples lacking preprocessed morphology images, maximum intensity projection (MIP) images were generated from 3D image stacks by taking the pixel-wise maximum across all z-planes. Focus images were generated by selecting the sharpest plane using Tenengrad focus measure computed on the full image plane). Images were rendered at multiple resolutions (100–1000 DPI) in both color and grayscale variants.

Hematoxylin and eosin (H&E) imaging data were available for 124 of 1,539 samples. Of these, 60 samples lacked the affine transformation matrix files, necessary for registering the H&E images to the morphology image coordinate space. We manually aligned these images using the image alignment tool in the 10x Genomics Xenium Explorer software.

### Annotation of Whole-Slide Images

Whole slide images (WSI) of breast cancer tissue slices were viewed by a surgical pathologist using Sedeen Pathcore Viewer. Slides were preliminarily screened for suitability for evaluation. Immunohistochemistry slides, redundant slides, and slides with poor quality were excluded. Before annotation, a histologic diagnosis was rendered based on criteria in the 5^th^ Edition of the WHO Classification of Breast Tumors (WHO Classification of Tumours Editorial Board 2019). Particular attention was given to foci of invasive carcinoma, nests of ductal carcinoma in situ, intratumoral fibrosis, entrapped adipocytes, and lymphovascular invasion. Whenever possible, annotation was made to individual foci of carcinoma and DCIS; however, in some instances, confluent foci of invasive carcinoma and DCIS were annotated as one group. Representative non-neoplastic glandular structures, i.e., terminal duct-lobular units, were also annotated for comparison. Users can inspect these annotations and additional comments in the Xenium interface.

### Database interface implementation

The database interface is built on top of the existing DeepSpaceDB database, with a backend implemented in Flask and a frontend using JavaScript with Plotly.js for interactive visualizations (Plotly Technologies Inc. 2015). While DeepSpaceDB has a Visium-oriented interface, this interface is not suitable for Xenium data. Xenium data often contains 100,000s to >1 million cells, compared to the typical 100s to 1,000s of spots in Visium data. Therefore, we developed a new Xenium-oriented interface with architectural optimizations to handle the increased data scale.

The interface selects the appropriate pre-computed data format based on whether the user is viewing or analyzing the data: gene-chunked stores are used to display individual genes, spatial-chunked stores are used for Region of Interest (ROI) analysis, and sparse matrix formats are used for single-cell resolution viewing mode. To keep data transfer minimal, the server removes cells or bins with zero expression before sending data to the client, which reduces the amount of data transmitted by 70-90%. During spatial plot rendering, cell coordinates are cached server-side, reducing transmitted data to label data such as gene expression values or cell-type predictions. When users analyze selected ROIs, all calculations are performed on the server using NumPy vectorized operations, and only the summary statistics are sent back. This reduces the time for ROI analysis and allows users to get their statistical results in less than a second for single-cell resolution selections. Statistical comparisons between regions use Welch’s t-test with Bonferroni correction for multiple testing.

On the client side, Plotly.js with WebGL acceleration handles efficient rendering of massive datasets. Point sizes automatically adjust as users zoom in and out to keep the visualization clear at any scale. When displaying multiple genes simultaneously, their expression patterns are combined using RGB color blending with colors weighted by expression intensity. Users can switch between different spatial resolutions or view different genes without reloading the entire dataset - the system preserves their analysis context (selected regions, zoom level, and interaction mode) when switching views. Caching of visualization settings is also implemented to avoid unnecessary requests to the server.

## RESULTS

### Overview of workflow and collected samples

We collected Xenium samples from NCBI GEO, the 10x Genomics website and DBKERO (Barrett *et al*. 2013, Suzuki *et al*. 2018), including samples covering smaller gene panels as well as Xenium Prime samples covering approximately 5 thousand genes. We developed a data processing pipeline that checks if the downloaded data is sufficient to correctly process each sample and then processes it into a SpatialData object (Figure 1 and Supplementary section “DeepSpaceDB Xenium Data Processing Pipeline”). Where needed, the pipeline converts data formats to be consistent with the input formats required for SpatialData. In total, 1,539 Xenium samples (including 213 million cells) were successfully processed and added to the DeepSpaceDB database. To facilitate access to this data from the database interface, each sample was pre-processed into a number of different formats (see Methods). In short, the single-cell resolution data of each sample was stored into a Zarr store chunked by gene, as well as a Zarr store that was chunked by spatial coordinates. In addition, data was binned into bins of various sizes (5, 10, 20, 40, 80 and 160 µm) and the resulting data was pre-processed into Zarr stores chunked by gene. Together, these different Zarr stores allow for short response times when using the viewing and analysis functions in DeepSpaceDB.

We conducted an analysis of the characteristics of the collected data. The median number of cells per sample was roughly 74k cells (range 103 to 2.5 million, mean 139k; Figure 2A), with a median of 12.6 million transcripts (range 1089 to 1.9 billion, mean 30.4 million; Figure 2B). The samples included 1,272 samples that use small gene panels (100∼500 genes) while 267 were Xenium Prime samples with roughly 5,000 genes. Naturally, a positive trend was observed between the number of cells and the number of transcripts per sample (Figure 2C) with Xenium Prime samples in general containing a higher number of cells and transcripts. The publication dates of the samples revealed that the number of samples being published is rapidly increasing (Supplementary Figure S1A). Overlap in covered genes was, in general, low; whereas each study tends to use the same gene panel for the samples it generates, different studies use different gene panels. No gene was covered in all samples (Supplementary Figure 1B-C) and only 121 genes are used in more than 50% of the human samples (162 genes for mouse). Following PCA analysis of several features of all samples, we also established a rough general quality score which was added to the database (see Supplementary section “Database-wide quality trends” and Supplementary Figure S2 and Table S1).

**Figure 2.**
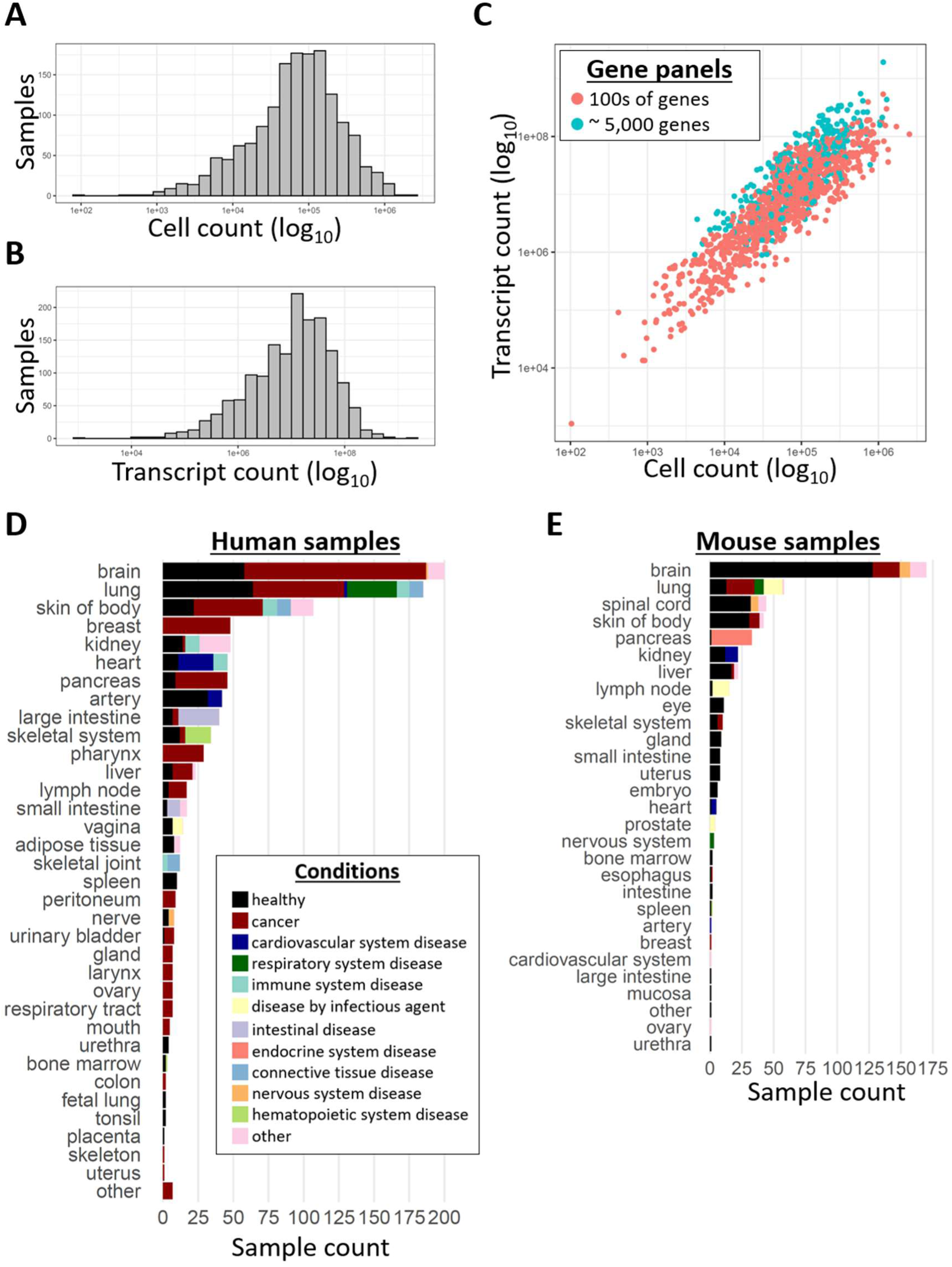
Characteristics of the 1,539 collected Xenium samples. **(A)** A histogram showing the number of cells per sample. **(B)** A histogram showing the total transcript count per sample. **(C)** A scatterplot showing the number of cells (x axis) versus the total transcript count (y axis) for all samples. Samples using smaller gene panels and samples covering roughly 5,000 genes are shown in red and blue, respectively. **(D-E)** The number of available samples for human (D) and mouse (E) per tissue. Colors indicate disease states.

Annotation data was manually collected for each processed sample, including, amongst others, the organ of origin, medical condition, publication date, related literature, etc. Samples included 992 human, 486 mouse, 29 axolotl (Ambystoma mexicanum), 8 little skate (Leucoraja erinaceus), 6 rat (Rattus norvegicus), and 1 lancelet (Branchiostoma floridae) samples, as well as 17 human organoids implanted into mice. Sample counts for human and mouse tissues are shown in Figure 2D,E. Human samples covered a wide variety of tissues and organs, and were especially rich in cancer-related samples. For mouse, brain samples were especially abundant.

### The DeepSpaceDB Xenium main interface

After selecting a Xenium sample in the DeepSpaceDB “Database” tab, the selected sample is shown in the Xenium interface. A brief overview of the main Xenium interface is shown in Figure 3A, using a human breast cancer example (sample ID DSIDX00001) as example (Janesick *et al*. 2023). The default view after selecting a sample includes sample information, a gene list, 5 submenus (“Panels”, “Export”, “Summary Statistics”, “Customize”, and “Compare Sample”), a tool bar, a Region Management menu (which appears after selecting ROIs), and – where available – a Xenium Analysis Summary. After selecting a sample, by default the cell clustering results are shown (Fig. 3A). The gene list includes all genes sorted by the strength of their spatial pattern, as judged by the singleCellHaystack method. Users can search for their gene of interest, or click one of the top scoring genes. The top-scoring gene in this sample is *FASN* (Fig. 3B). If needed, the image data can be hidden, as illustrated for *CEACAM6* (Figure 3C), which has a distinct spatial pattern from *FASN*. When selecting a gene to plot, the database employs the pre-calculated Zarr stores that have been chunked by gene to quickly visualize the selected gene. When the “Multi-Gene Mode” is turned on, users can plot the expression of multiple genes in 1 plot. An example plotting both *FASN* and *CEACAM6* is shown in Figure 3D. Users can plot a variety of features including clustering results in the spatial coordinates (Fig. 3A) or in a 2D embedding (Supplementary Figure S3A). Spatial domains predicted by BANKSY can also be inspected (Fig. 3E and Supplementary Figure S3B). In addition, cell type predictions (Fig. 3F) and quality measures could be plotted, such as in the number of detected genes per cell (Figure 3G). For a subset of human breast cancer samples, pathology image annotations are also available (Fig. 3H). Users can also plot cell and nucleus boundaries when available (Supplementary Figure S3C). Finally, in addition to plotting the data in single-cell resolution, users can also choose to plot the data processed to various bin sizes (ex: bin size 40 µm in Supplementary Figure S3D) or individual transcripts (Supplementary Figure S3E).

**Figure 3.**
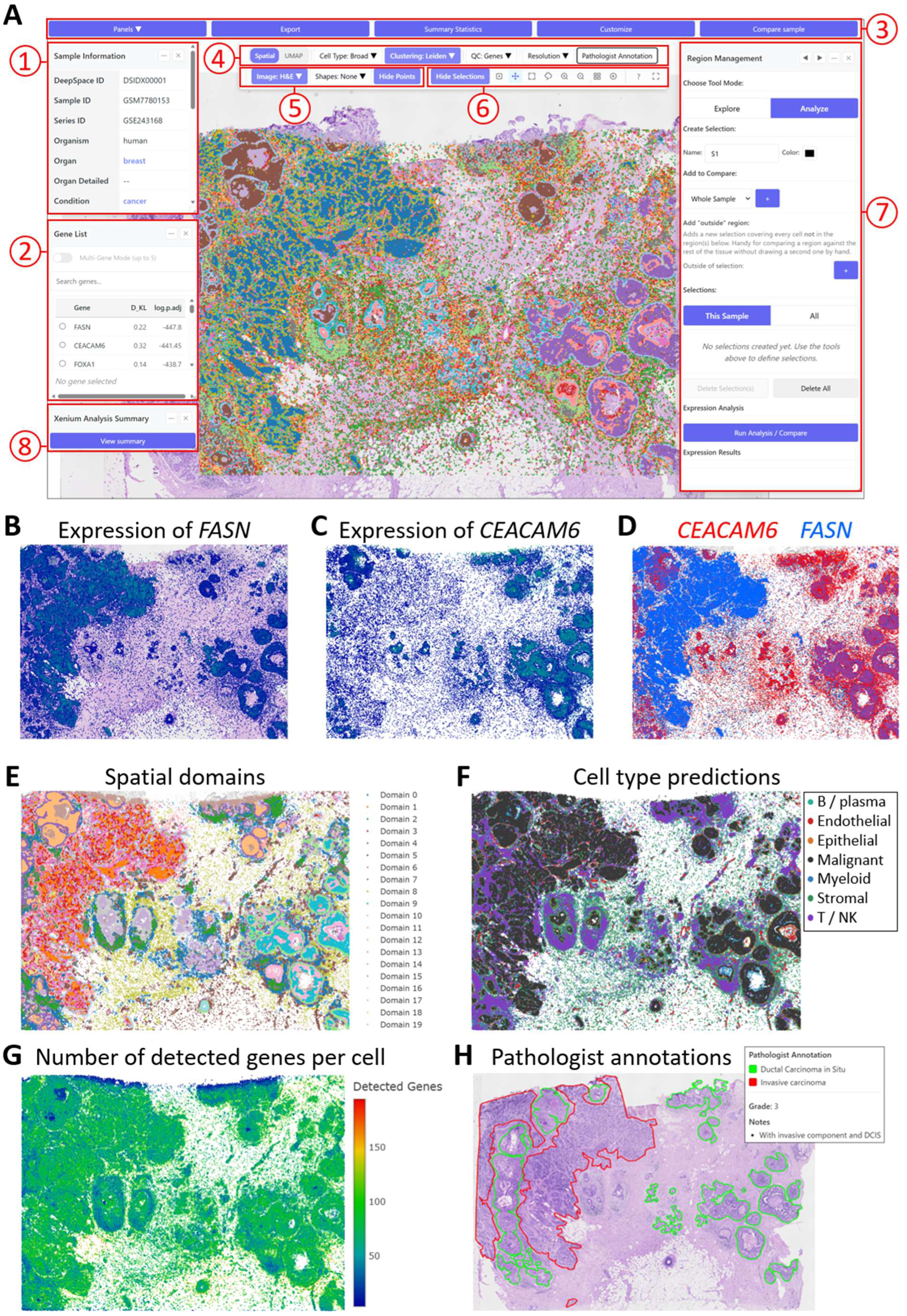
Overview of the DeepSpaceDB Xenium interface. **(A)** The main sample view of the Xenium interface. The cell clustering result of a human breast cancer sample (sample ID DSIDX00001) is shown on top of the H&E image of this sample. Indicated are 1: sample information. 2: A list of genes sorted by the strength of their spatial pattern. 3: Submenus “Panels”, “Export”, “Summary Statistics”, “Customize”, and “Compare Samples”. 4: A tool bar controlling what is plotted. 5: A toolbar where users can toggle on/off images, cells and boundaries. 6: A toolbar for panning, selecting, and zooming in/out. 7: The region management tool where selected regions are shown and can be compared (see also Fig. 4A). 8: A link to the Xenium Analysis Summary of this sample. **(B)** The expression of *FASN* with the H&E image in the background. **(C)** The expression of *CEACAM6* is plotted without H&E image in the background. **(D)** The expression pattern of both *FASN* (blue) and *CEACAM6* (red) in one plot. **(E)** Predicted spatial domains. **(F)** Predicted cell types. **(G)** The number of detected genes per cell. **(H)** H&E image annotations for this tissue slice.

The “Export” submenu offers options for downloading the Xenium data and exporting plots. Users can download both the raw data collected from the original source (GEO, 10x Genomics website or DBKERO), as well as the data files produced by our pipeline. If the data has no issues, the files produced by our pipeline are identical to the raw data, apart from having standardized names that SpatialData package requires. Otherwise, we provide cleaned and salvaged raw files suitable for loading using Python or R, or desktop viewers such as Xenium Explorer. Users can also select a subset of data files to download. The “Customize” submenu allows users to change the aesthetics of the plotted data, including color scales and the image to plot (MIP, focus, or H&E where available), as well as export the plot as a raster or vector image.

### Interactive investigation of a region of interest

In the tool bar, users can hide or view the image data and the transcriptome data. There are also tools to zoom in/out and to select ROIs. ROIs can be selected by using the mouse cursor (using a box or lasso) or by selecting domains, clusters, cell types, or pathology annotations. As an example, we here use pathology annotations to select one of the large invasive carcinoma regions (region 15 in the sample; Figure 4A). After clicking the “Run Analysis / Compare” button, several analysis functions become available. One allows users to investigate gene co-expression patterns within the selected ROI. For example, several genes have a high positive or negative correlation of expression with *FASN* (Figure 4B). By plotting the expression of both *FASN* and *LUM* (the gene with the highest negative correlation with *FASN*) in the same plot we can confirm that these two genes tend to be expressed in different cells (Figure 4C). Another function shows the predicted cell type composition of the selected ROI (Figure 4D), confirming that this region is composed mainly of malignant cells. Spatial neighborhood enrichment analysis can be used to analyze whether pairs of cell types tend to be located in proximity to each other within the selected ROI. Figure 4E reveals that cells of the same cell type tend to be in general located together (positive Z-scores on the diagonal). This is especially the case for malignant cells (highest Z-score). Indeed, the ROI contains a large number of aggregates of malignant cells (Figure 4F shows one representative region). The heatmap also reveals that malignant cells tend to be less located in proximity to other cell types (negative Z-scores). Among pairs of different cell types, myeloid cells and T / NK cells have the highest tendency for co-localization. This too, can be visually confirmed (Figure 4F).

**Figure 4.**
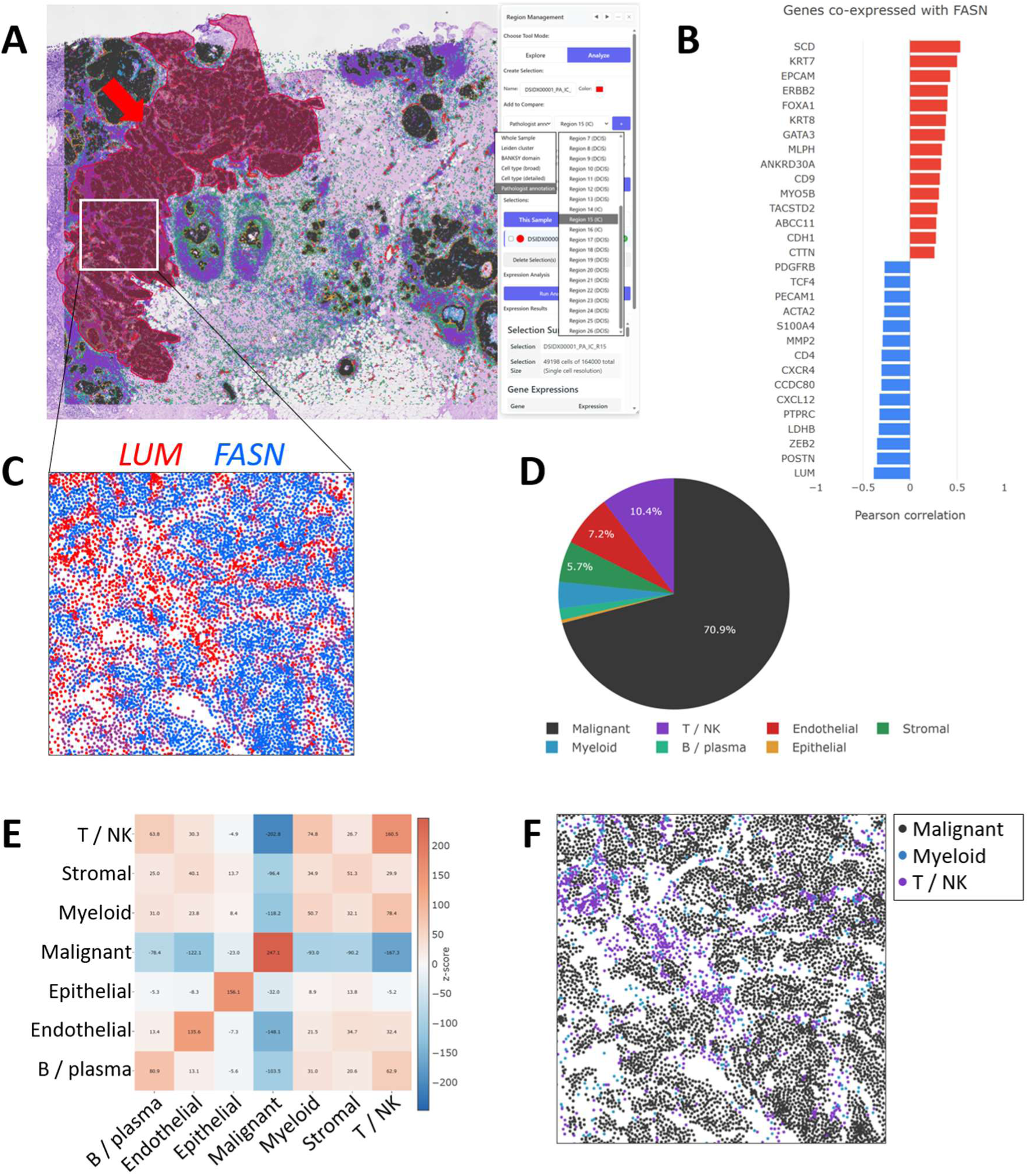
Interactive exploration of a single ROI using the DeepSpaceDB Xenium interface. **(A)** A view on the sample shown in Figure 3A, with a large invasive carcinoma region of interest (ROI) selected (indicated in red and by the red arrow). **(B)** Genes with high positive and negative correlation of expression with FASN within the selected ROI. **(C)** The expression patterns of FASN (in blue) and LUM (in red) in a part of the ROI. LUM has the highest anti-correlation with FASN in the ROI. **(D)** A pie chart showing the predicted cell type composition of the selected ROI. **(E)** Heatmap summarizing spatial neighborhood enrichment analysis within the selected ROI. **(F)** The spatial distribution of malignant, myeloid, and T/NK cells in a part of the ROI. For clarity, other cell types are not plotted.

### Comparison between multiple regions of interest

Users can select multiple ROIs and compare between them. As an example, we selected two ROIs in the breast cancer slice (Figure 5A) using the lasso tool, one overlapping with ductal carcinoma in situ (DCIS) tissue (selection 1) and one outside of the DCIS region (selection 2). When one or more regions have been selected, the “Region Management” menu appears (Figure 5A, menu on the right). When the user clicks “Analyze” and “Run Analysis / Compare”, the average expression of all genes in each ROI is calculated. The result is available for download and can be visualized using volcano plots and scatter plots (Figure 5B and Supplementary Figure S4A). Hovering above points in the plots shows information including the gene symbol, which can subsequently be used to confirm the spatial expression patterns (Figure 5C). For example, genes *KRT8*, *MLPH*, and *EPCAM* have a higher expression in the 1^st^ selected region, while *CXCL12*, *LUM*, and *PTGDS* have a higher expression in the 2^nd^ region. The selection of the ROIs, generation of the plots, and confirmation of differentially expressed genes can be done in seconds using this interface. As a result of the client and server–side optimizations, our viewer allows fast and efficient access to the Xenium data without the need to download it, surpassing the speed of any other available web-viewers.

**Figure 5.**
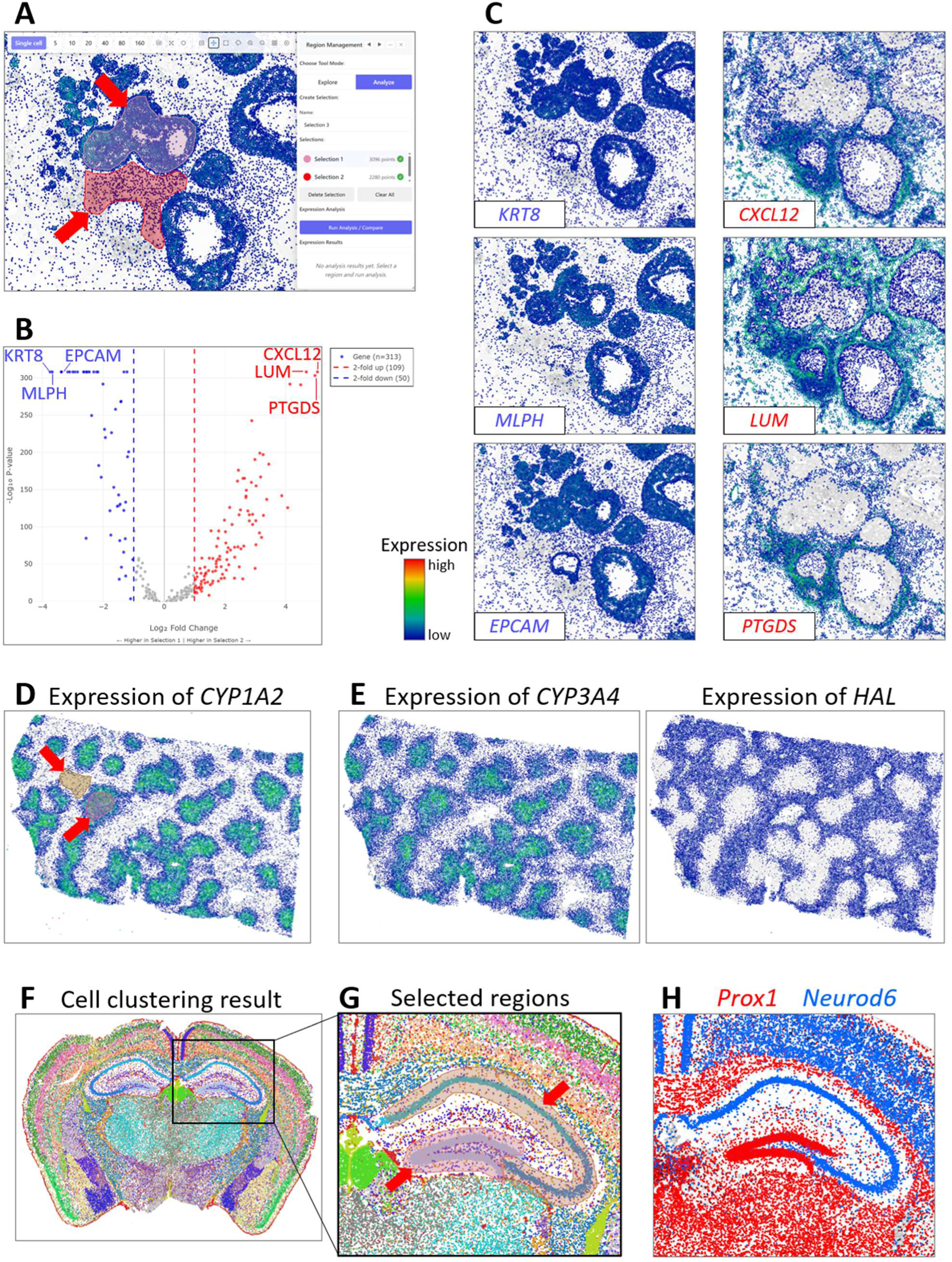
Interactive exploration of multiple ROIs using the DeepSpaceDB Xenium interface. **(A)** A zoomed-in view on the sample shown in Figure 3A, with 2 selected regions of interest (ROIs, indicated by the red arrows). **(B)** A volcano plot of the gene expression levels in the 2 selected ROIs. Genes with large differences in expression are indicated. **(C)** The expression patterns of the 6 indicated genes, three of which have a high expression in ROI1 (left side) and three with a high expression in ROI2 (right side). **(D)** The expression of gene *CYP1A2* in a human liver tissue (sample ID DSIDX00197) is used to select 2 ROIs (*CYP1A2*^high^ and *CYP1A2*^low^). **(E)** The expression patterns of two genes with differential expression between the two ROIs. *CYP3A4* and *HAL* have high expression in the *CYP1A2*^high^ and *CYP1A2*^low^ regions, respectively. **(F)** The cell clustering result of a mouse brain sample (sample ID DSIDX00160). **(G)** A zoomed-in view on the hippocampal region of the mouse brain tissue, with two selected ROIs, roughly corresponding to the dentate gyrus and Ammon’s horn. **(H)** The expression pattern of two differentially expressed genes in 1 plot. Prox1 (red) and *Neurod6* (blue) have high expression in the dentate gyrus and Ammon’s horn, respectively.

### Usage examples

Searching DeepSpaceDB for human liver samples of the Xenium platform returns 9 hits. Selecting sample DSIDX00197, we can confirm the zonated expression pattern of gene CYP1A2, with high expression in the pericentral regions of the liver (Figure 5D). To identify other genes with zonated expression patterns, we selected ROIs in a CYP1A2^high^ area (pericentral) and a CYP1A2^low^ area (periportal) and compared expression between these 2 regions (Supplementary Figure S4B). Figure 5E shows 2 of the high-scoring genes of each zone (*CYP3A4* for the pericentral and *HAL* for the periportal zones, respectively). Finally, we use the Xenium interface to explore differential expression in the mouse brain. By inspecting the clustering result of sample DSIDX00160 (Figure 5F), we can clearly observe known structures of the brain, including layers of the cortex, the thalamus and hypothalamus. Using the interface to zoom in on the hippocampus, we select 2 ROIs, one roughly consisting of the dentate gyrus and one of Ammon’s horn (Figure 5G). Through comparison of the gene expression between these 2 regions we can identify several genes with significantly differential expression (Supplementary Figure S4C). Two of them are plotted together in Figure 5H (*Prox1* in red with high expression in the dentate gyrus and *Neurod6* in blue with high expression in Ammon’s horn). Thus, in a matter of seconds, the DeepSpaceDB Xenium interface can be used to explore liver zonation-associated genes or differentially expressed genes within substructures of various tissues.

### Inter-sample comparisons

Users are not restricted to comparisons between ROIs within the same sample; inter-sample comparisons are also possible, through the “Compare Sample” menu (see Fig. 3A). However, it should be noted that such comparisons are limited by batch effects and by the fact that different studies tend to use different gene panels and might therefore have only few genes in common (Atta *et al*. 2024). Here, we present one inter-sample comparison using two breast cancer samples from two different patients, generated by the same study using the same gene panel (Janesick *et al*. 2023). We selected all invasive carcinoma regions in both samples (3 in sample DSIDX00001 and 14 in DSIDX00003; Figure 6A) and predicted DEGs between them (Figure 6B). Among the genes with higher expression in the invasive tumor regions of sample DSIDX00001 and DSIDX00003 were *GATA3* and *S100A8*, respectively. Visual inspection confirms these findings (Figure 6C-D). Moreover, sample DSIDX00001 is T2N1M0, Stage II-B, ER+/PR−/HER2+, whereas DSIDX00003 is pT2 pN1a pMX, ER−/PR−/HER2+ (Janesick *et al*. 2023). The higher *GATA3* signal in DSIDX00001 is consistent with its ER-positive status, whereas *S100A8* likely reflects differences in the myeloid component. This comparison therefore captures both tumour-cell-intrinsic differences and differences in immune composition.

**Figure 6.**
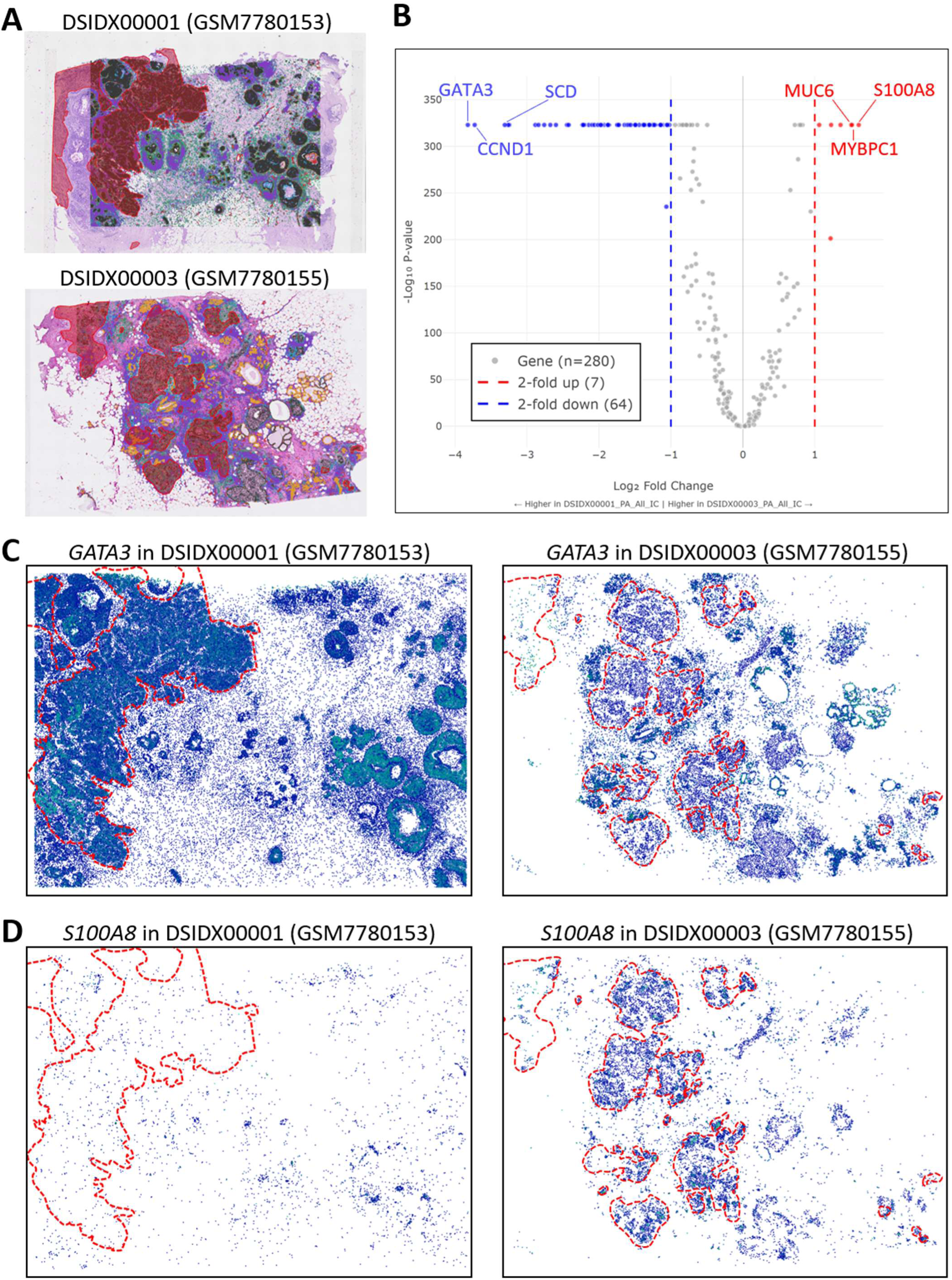
Conducting a comparison between two samples. **(A)** The two breast cancer samples used in this comparison. In each, invasive carcinoma regions have been selected for this comparison (3 in the top sample and 14 in the bottom sample; indicated in red). **(B)** A volcano plot of the gene expression levels in the selected ROIs. Genes with large differences in expression are indicated. **(C-D)** The expression patterns of GATA3 and S100A8 in both samples. Expression of GATA3 is higher in the invasive carcinoma regions of sample DSIDX00001 (left), and expression of S100A8 is higher in the invasive carcinoma regions of sample DSIDX00003 (right). For clarity the selected ROIs are indicated by dotted red lines and the tissue images are not shown.

### Searching the DeepSpaceDB for genes with spatial expression patterns

In addition to the analysis of individual samples, DeepSpaceDB allows users to query the entire collection for a gene of interest. This search function was introduced for the Visium samples in DeepSpaceDB (Honcharuk et al. 2025) and has now been extended to the Xenium collection. Under the “Search” tab, a user selects a species and platform and enters a query gene. All samples containing that gene are returned, ordered by the strength of its spatial expression pattern as measured by singleCellHaystack, with links to the original data source and to the sample in the Xenium interface, as well as a preview of the sample image and of the expression of the queried gene. Since these values are pre-computed, results are returned instantly. DeepSpaceDB additionally tests whether samples of a given tissue or medical condition are enriched among the highest-ranking samples, using a one-sided Mann-Whitney U test with correction for multiple testing.

As an example, the results of searching the human Xenium samples for CEACAM6, EPCAM, FOXP3, and EGFR are summarized in Figure 7. For CEACAM6, samples with strong spatial patterns are enriched for large intestine samples and patients suffering from intestinal diseases (Figure 7A). The top scoring sample (DSIDX00398) is a colon cancer sample and shows a clear spatial pattern of expression for CEACAM6. Similarly, EPCAM shows spatially variable expression patterns in cancer-related samples, including breast cancer samples (Figure 7B), with the top-scoring sample being a colon cancer sample. FOXP3 has strong spatial patterns in lymph node samples and cancer samples, with the strongest pattern being found in a tonsil sample (Figure 7C). Finally, cancer-related samples have clear patterns of expression of EGFR, including a brain glioblastoma sample (Figure 7D). In this way, the Search tool can be used to query the entire DeepSpaceDB database and quickly find samples with clear patterns of expression of a gene of interest.

**Figure 7.**
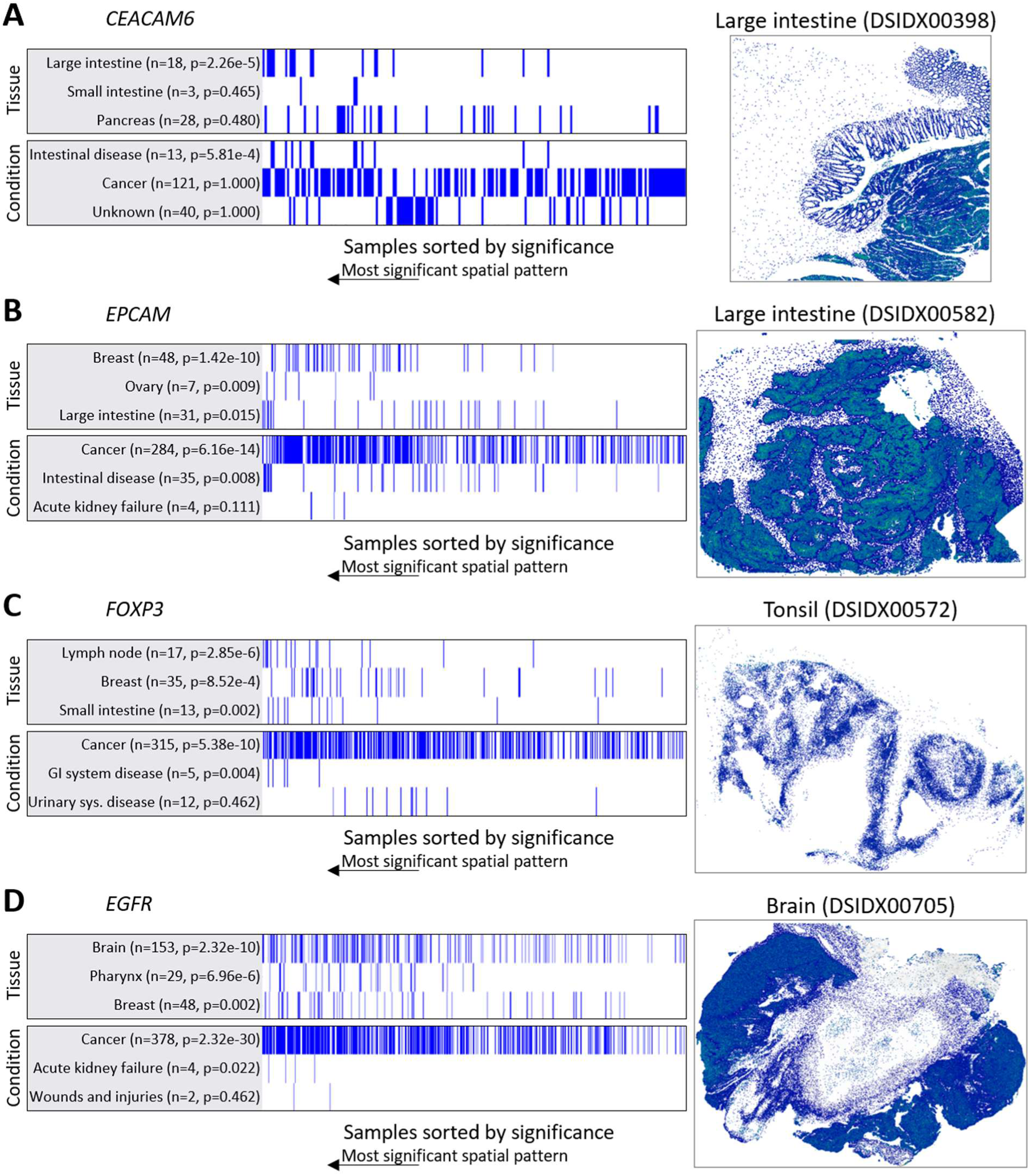
Searching DeepSpaceDB Xenium data with a query gene. For each query a ranking of all samples sorted by significance of the spatial pattern (as judged by the singleCellHaystack method) of the query gene is shown on the left. **(A)** CEACAM6. **(B)** EPCAM. **(C)** FOXP3. **(D)** EGFR. For each gene, tissues and conditions are shown, along with the number of samples (n) and the p-value of a one-sided Mann-Whitney U test (p), adjusted for multiple testing. Tissues or conditions are sorted by the p-value. Only the top 3 tissues and conditions are shown. At the right hand side, the top-scoring sample (i.e., the sample with strongest spatial pattern for the query gene) is shown with the expression of the query gene.

## DISCUSSION

The rapid accumulation of public high-resolution spatial transcriptomics data presents both an opportunity and a challenge for the scientific community. While Xenium enables single-cell resolution measurements across large tissue sections, the reuse of these data is hindered by their size, complexity, and inconsistent public release formats. In this work, we addressed this challenge by developing a data processing pipeline and a new interactive web-based interface. Together, these enable fast exploration of public Xenium datasets.

A major obstacle to reusing public Xenium data is the lack of uniformity in released files, particularly for datasets deposited in general-purpose repositories such as GEO. Many samples deviate from the canonical Xenium Analyzer output structure, and a substantial fraction is incomplete. By implementing a validation and recovery pipeline that detects, repairs, and, when possible, regenerates missing components, we were able to successfully process and integrate a large proportion of collected samples. Clearer instructions by companies providing these platforms and by the data archives about the minimal necessary files needed for reanalysis would greatly facilitate reproducibility of spatial genomics data analysis.

The scale and access patterns of Xenium data further necessitate careful consideration of data representation. Gene-centric queries, such as visualizing the spatial distribution of a single gene, and spatially localized queries, such as extracting expression profiles from regions of interest, impose fundamentally different computational requirements. Our use of complementary gene-chunked and spatially chunked Zarr stores explicitly addresses these differing access patterns and avoids expensive reshaping operations at query time. Together with multi-resolution spatial binning, this design enables sub-second response times for common interactive operations, even for samples containing hundreds of thousands to more than a million cells. These architectural choices are critical for maintaining interactivity in a web-based environment and would be difficult to achieve using a single, monolithic data representation, making DeepSpaceDB Xenium interface considerably faster than any other existing web-based solutions.

An important observation from our analysis of collected datasets is the limited overlap in gene panels across studies, with no single gene present in all samples. This fragmentation poses a fundamental limitation for cross-study analyses. Previously, we were able to merge all collected Visium samples into a global atlas covering millions of spots, and use this merged atlas as a basis for spot-level annotation (Honcharuk et al. 2025). A similar integrative analysis is difficult for large collections of Xenium samples because very few genes are covered in a large subset of all samples. Nevertheless, comparisons between samples generated within the same study with similar gene panels are possible (Figure 6).

Several tools exist for visualizing Xenium data, including proprietary solutions such as Xenium Explorer and local visualization workflows based on napari (napari contributors 2019). Proprietary viewers offer high performance but are restricted to selected datasets and do not support unified access to public collections. Local tools, while flexible, require full data download and substantial computational resources, limiting their accessibility. The DeepSpaceDB Xenium interface complements these approaches by providing fast, browser-based access to a large collection of public datasets without requiring local data storage or specialized hardware. Users requiring advanced or customized analyses can download processed data from DeepSpaceDB and continue their work in dedicated analysis environments.

This work has several limitations. The quality and completeness of metadata are constrained by what is reported in original studies, and despite extensive preprocessing, inherited biases and inconsistencies remain. Statistical analyses performed within the interface prioritize speed and interactivity over methodological complexity and are not intended to replace comprehensive downstream analyses. In addition, limited gene panel overlap restricts certain types of cross-sample comparisons. Furthermore, to ensure smooth exploration using the interface, single cells are represented as points by default rather than as polygons, and images are shown in a relatively low resolution. However, cell and nucleus boundaries can be displayed when needed. For analyses requiring detailed cell shapes and high-resolution image data, users can download the data files provided on DeepSpaceDB and conduct a more thorough analysis locally. Finally, while cell type predictions are assigned to cell clusters, no anatomical labels have been given to the predicted spatial domains.

DeepSpaceDB is updated regularly with newly released samples and with new analysis methods. Future development will focus on addressing the limited overlap between gene panels, which is the main obstacle to broader cross-dataset analysis. One approach is to describe cells not by individual genes but by scores over predefined gene sets, such as Hallmark and Reactome pathways or published cell type signatures (Liberzon et al. 2015, Ragueneau et al. 2026). Such scores can be estimated as long as a sufficient fraction of the genes in a set is present on a panel, providing a feature space that is shared between samples with different panels, at the cost of gene-level resolution. Applying the existing inter-sample comparison function of DeepSpaceDB in such a space would extend it to comparisons across studies, tissues, and disease conditions.

## DATA AVAILABILITY

Data of all samples can be downloaded from the DeepSpaceDB website. Links to the original data source are also provided for each sample.

## CODE AVAILABILITY

The Xenium data processing pipeline is available in a GitHub repository (https://github.com/vladyslav-honcharuk/deepspacedb-xenium-pipeline).

## FUNDING

This work was supported by the Japan Science and Technology (JST) Agency National Bioscience Database Center (NBDC) [JPMJND2303 to A.V.], JST FOREST [JPMJFR2062 to K.S.], JST Moonshot [JPMJMS2011-61 to K.S.], by the Japan Agency for Medical Research and Development [JP24gm2010003 to A.V.], by the Japan Society for the Promotion of Science (JSPS) [JP26HP8021 to A.V., JP24K02236 to K.S., JP24KK0147 to K.S., and JP25K21774 to K.S.], by MEXT Promotion of Development of a Joint Usage / Research System Project: Coalition of Universities for Research Excellence Program (CURE) [JPMXP1323015484 to K.S.]. and by an Office of Directors’ Research Grant provided by the Institute for Life and Medical Sciences of Kyoto University (to A.V.). The funders had no role in study design, data collection and analysis, decision to publish, or preparation of the manuscript.

## CONFLICT OF INTEREST

The authors declare no conflict of interest exists in this study.

## ACKNOWLEDGEMENTS

The authors would like to thank Y. Harada for secretarial assistance.

## AUTHOR CONTRIBUTIONS

Alexis Vandenbon: Conceptualization, Formal analysis, Software, Writing – original draft, Writing – review & editing, Funding acquisition. Vladyslav Honcharuk: Software, Data curation, Formal analysis. Keiko Takemoto: Software, Data curation. Marvin C. Masalunga: Formal analysis. Hongyi Zhao: Formal analysis. Diego Diez: Formal analysis, Software. Shinpei Kawaoka: Formal analysis, Software. All authors: Writing – review & editing.

## Notes

http://www.deepspacedb.com/

### Competing Interest Statement

The authors have declared no competing interest.

### Summary of Updates

List of the main changes: 1.The number of integrated Xenium samples has increased from 628 to 1,539, collected from 1,831 initially retrieved samples, and the collection now covers six species. 2.Cell type predictions have been added for all human and mouse samples, based on reference profiles derived from CellxGene, using the same set of cell type labels across all samples (new Methods subsection; Figure 3F). 3.Spatial neighborhood enrichment analysis, gene co-expression analysis, and cell type composition analysis of user-selected regions of interest have been added (new Methods subsection; Figure 4B, D, E). 4.A "Compare Sample" function has been added, allowing comparison of gene expression between selections of cells in two different samples (new Results subsection "Inter-sample comparisons"; Figure 6). 5.The database-wide gene search function, previously available for Visium samples only, has been extended to the Xenium collection (new Results subsection; Figure 7). 6.Cell and nucleus boundaries and individual transcript locations can now be displayed in the interface (Supplementary Figure S3C, S3E). 7.Expert pathology annotations of H&E images have been added for a subset of human breast cancer samples, and can be used directly as regions of interest (Figure 3H). 8.The data processing pipeline has been made publicly available, and a Code Availability statement has been added. A new Supplementary section describing the validation and repair procedures applied to heterogeneous and incomplete Xenium inputs has been added. 9.The final paragraph of the Discussion has been rewritten to describe planned developments concretely, and the limitations paragraph has been extended.

