## Supplementary Material for "DeepSpaceDB 2.0: an interactive spatial transcriptomics database for large-scale Xenium data exploration"

### SUPPLEMENTARY METHODS

#### DeepSpaceDB Xenium Data Processing Pipeline

To collect and process all publicly available 10x Genomics Xenium data from the NCBI Gene Expression Omnibus (GEO), we developed a standardized pipeline capable of identifying Xenium sample records, downloading their associated data, resolving data inconsistencies and standardizing the datasets by transforming them into SpatialData (Marconato *et al.* 2025) objects followed by serialization.

The set of files produced by the Xenium Onboard Analysis software follows a defined specification, while the SpatialData package expects specific files and filenames to be preserved. The standard Xenium output includes an experiment manifest file (*experiment.xenium*), morphology images (*morphology.ome.tif*, *morphology_mip.ome.tif,* and *morphology_focus.ome.tif*), cell-level information (*cells.csv.gz* or *cells.parquet*), cell and nucleus segmentation boundaries (*cell_boundaries.parquet* and *nucleus_boundaries.parquet*), segmentation masks (*cells.zarr.zip*), transcript-level data (*transcripts.parquet* or *transcripts.zarr.zip*), and cell-feature matrices (*cell_feature_matrix/*, *cell_feature_matrix.h5*, or *cell_feature_matrix.zarr.zip*), together with additional metadata and analysis outputs (see https://www.10xgenomics.com/support/software/xenium-onboard-analysis/latest/analysis/xoa-output-at-a-glance).

Although NCBI GEO provides recommendations regarding the files that should be submitted, these recommendations do not encompass the complete set of files required for downstream analyses. We found that the majority of GEO submissions deviate from the output format produced by the Xenium software suite and expected by SpatialData, with substantial heterogeneity in both file formats and filenames. A substantial proportion of submissions also lack files required for reconstructing complete spatial datasets, rendering them incompatible with standard downstream processing. Therefore, we developed a pipeline capable of processing raw uploaded data and converting it into a standardized SpatialData object (https://github.com/vladyslav-honcharuk/deepspacedb-xenium-pipeline). The pipeline incorporates a range of recovery procedures and workarounds to resolve inconsistencies, reconstruct missing or non-standardized file structures where possible, and extract usable information from incomplete submissions.

The pipeline consists of four main stages: (1) identifying all Xenium samples in GEO, (2) downloading their associated data, (3) verifying the presence of the necessary files and recovering usable data when submissions deviate from the standard file formats and naming conventions, and (4) constructing a standardized SpatialData object from the processed data. This enables the automated processing of GEO Xenium submissions into standardized SpatialData objects without manual intervention.

(1) Identifying Xenium samples in GEO

We queried the NCBI E-utilities API (eutilities.github.io) against the GEO DataSets database (db=gds). The pipeline first runs ESearch using the query "Xenium[All Fields] AND GPL" and retains UIDs beginning with 1, corresponding to GEO platform (GPL) records. The ESummary is then used to retrieve the corresponding platform accessions. Each individual platform contains submissions for an individual species.

For each identified platform, the pipeline queries "{GPL}[ACCN]" and retains UIDs beginning with 3, corresponding to GEO sample (GSM) records. Each GSM is cross-referenced with its associated GSE using the linked series information in the same summary record. This produces a platform–series–sample index of identified Xenium samples.

(2) Downloading Xenium associated data

For each GSM, the pipeline resolves the corresponding GEO MiniML record (geometa.cgi?mode=miniml) to retrieve supplementary-data file URLs and the associated GPL and GSE identifiers from the Platform-Ref and Series-Ref fields. All supplementary files are downloaded into a standardized directory structure: {GPL}/{GSE}/{GSM}/raw/. A ".download_complete" marker file is generated following successful completion, allowing interrupted downloads to be resumed without reprocessing completed samples.

(3) Verification and recovery of data files

The main stage of the pipeline is data salvage and standardization. The pipeline first determines whether the submitted data consists of individual files or an archive. For archived submissions, the archive format is identified (i.e.: .tar.gz, .zip, .gz, or .zst) and its contents are extracted. Once individual files are available, the pipeline verifies the presence and format of the required Xenium files and checks whether they follow the expected naming conventions. When files are missing, mislabeled, incorrectly encoded, or otherwise inconsistent with the expected structure, the pipeline attempts to recover and standardize the available data before downstream processing.

The most common inconsistencies identified in GEO submissions and the corresponding recovery procedures implemented by the pipeline are described below.

- *Mislabeled compression.* Files with a .gz extension are validated for gzip encoding. Files containing raw zlib/DEFLATE streams or uncompressed data are detected and re-encoded as valid gzip files.
- *Mislabeled file formats.* Files with extensions indicating a specific format are validated against their actual contents. For example, files named .parquet that cannot be parsed as Parquet are tested as CSV and, when valid, converted to correctly encoded Parquet files.
- *Non-standard filenames.* Arbitrarily named files are mapped to canonical Xenium filenames using an ordered, case-insensitive matching table, with the first matching rule applied. The mapping covers essential Xenium files, including cell-feature matrices, transcripts, cell and nucleus boundaries, morphology images, and gene-panel information.
- *H&E image identification.* H&E images are identified using token-aware regular expressions based on word boundaries and camel-case patterns rather than simple substring matching. This prevents false matches for filenames containing the token "he" within unrelated terms.
- *Reconstruction of experiment.xenium.* When *experiment.xenium* is absent, it is reconstructed from a complete template. Relevant metadata, including pixel size, panel,

instrument, and UUID fields, are populated where available. Image and explorer file references are subsequently updated after the corresponding output files have been generated.

- *Reconstruction of cells.parquet.* When *cells.parquet* is missing or malformed, it is regenerated with an enforced column order and predefined data types to ensure compatibility with downstream processing.
- *H&E-to-Xenium image alignment*. The pipeline searches for alignment or homography matrices using multiple known naming conventions. When alignment information is available, a scikit-image-based affine registration is applied to generate a pyramidal, aligned OME-TIFF in the Xenium pixel coordinate system.
- *Morphology image reconciliation.* Multi-page or multi-Z morphology stacks are converted into maximum-intensity projections. For files larger than 5 GB, pages are processed sequentially to avoid loading the entire stack into memory. The sharpest Z-plane is selected using a Tenengrad/Sobel focus measure (Piao *et al.* 2025), and multi-resolution pyramidal OME-TIFF files are generated. Morphology focus data provided as a single file, a directory of individual planes, or a tar archive are also normalized into the file layout expected by downstream processing.
- *Segmentation format compatibility.* The pipeline examines cells.zarr.zip for the mask and polygon keys required by the reader. The recorded *analysis_sw_version* is also used to distinguish pre- and post-2.0 segmentation formats, allowing incompatible labels from re-segmented datasets to be excluded when loading them would otherwise cause the standard reader to fail.
- *Expression matrix reconstruction.* As a last resort, when no pre-built expression matrix is available, the pipeline reconstructs a cell-by-gene sparse count matrix directly from transcripts.parquet. Negative-control, blank, deprecated, and unassigned codewords are excluded based on predefined name patterns.

(4) Construction of standardized SpatialData objects

The final stage constructs a standardized SpatialData object. The pipeline first attempts to use the standard spatialdata_io Xenium reader with capability flags configured according to the files available for each sample. If this fails, the pipeline falls back to a legacy pre-v3 HDF5-

based loading path and subsequently attempts four progressively simplified image configurations before classifying the sample as unsuccessful.

Following successful object construction, the pipeline performs standardized downstream analyses, including quality-control filtering and metric calculation, total-count normalization, log1p transformation, PCA, neighborhood graph construction, UMAP, and Leiden clustering (see Methods section of the main paper). Raw, normalized, and log1p-transformed expression values are retained as separate layers. Image rendering, correction-factor processing, and cell-shape overlays are also generated.

For efficient interactive visualization, expression data are spatially binned at multiple configurable resolutions and stored as chunked Zarr arrays. The complete single-cell sparse expression matrix is additionally exported in three layouts (CSR, CSC, and per-gene chunked representations), together with raw spatial coordinates. Binning and single-cell export are performed in parallel.

Each processing stage writes an independent completion or failure marker. Consequently, interrupted runs can be resumed, and subsequent executions reprocess only samples or stages that remain incomplete or have previously failed. At the end of each batch, the pipeline generates a QC summary .csv file and a list of successfully processed samples.

The heterogeneity observed in GEO submissions reflects differences in laboratory export procedures, software versions, and manual curation. In practice, submissions may contain files with the correct extension but incompatible encoding (e.g., .gz files that are not gzip-compressed or .parquet files containing CSV data), morphology focus stacks provided as a single tar archive, a directory of TIFF files, or a single TIFF, and missing or corrupted experiment.xenium and cells.parquet files.

Different versions of Xenium analysis software may also produce segmentation label formats that are not interchangeable. In many cases, deposits contain transcript-level data without a pre-computed expression matrix. These variations necessitate normalization before standardized downstream processing can be performed.

Each of these variations may result from differences in laboratory export procedures, software versions, or manual curation practices. Consequently, downstream processing cannot reliably assume a fixed file layout across GEO Xenium submissions without first applying a normalization step.

SpatialData construction through spatialdata_io.xenium expects a defined set of files and internal data structures. Depending on the available modalities, these include *cell_feature_matrix.h5/.zarr.zip*, *transcripts.parquet*, *cells.parquet/cells.zarr.zip*, *cell_boundaries.parquet*, *nucleus_boundaries.parquet*, *experiment.xenium*, *morphology_mip.ome.tif*, *morphology_focus.ome.tif* or a corresponding *morphology_focus/* directory, and *he_image.ome.tif*. The reader also relies on structural information within these files, such as specific keys in *cells.zarr.zip* and metadata fields in *experiment.xenium*, to construct tables, shapes, points, images, and coordinate transformations. SpatialData package simply fails to process the data that is not following the naming conventions or lacks even one file, such as lack of *experiment.xenium* file.

Consequently, deviations in filenames, internal encodings, or required metadata can prevent individual modalities from being loaded or cause failures during object construction. The mapping of heterogeneous GEO submissions to the standardized SpatialData schema therefore requires explicit normalization and validation rather than simple file renaming.

The pipeline converts an otherwise manual and error-prone curation process, including inspection of individual laboratory submissions, recovery of malformed files, mapping of heterogeneous filenames, and manual execution of downstream analyses, into a reproducible, resumable, batch-scale workflow. It enables automated discovery, retrieval, normalization, and analysis of publicly available Xenium GEO samples using a common processing framework.

By producing SpatialData objects with consistently structured QC metrics, clustering results, imaging data, spatial binning, and single-cell exports, the pipeline enables DeepSpaceDB to provide standardized cross-dataset interactive analysis without requiring manual curation of each individual sample.

Other observed trends in Xenium data submitted to GEO

We observed that more recent GEO submissions generally contained fewer structural inconsistencies, whereas earlier submissions exhibited greater heterogeneity. This trend coincided with the adoption of more stable versions of Xenium analysis software; however, the contribution of software version relative to other factors cannot be determined from the submissions alone.

An additional source of information loss was the selective deposition of Xenium outputs. 10x Genomics identifies decoded transcripts and high-resolution morphology images as core archival outputs for future reanalysis, while other Xenium outputs are derived from these data (https://www.10xgenomics.com/support/software/xenium-onboard-analysis/latest/analysis/xoa-output-archive-data). However, transcripts and morphology images alone do not constitute the complete Xenium output bundle required by downstream software such as Xenium Ranger. Reconstructing the derived outputs from these files requires additional analysis, including cell segmentation and transcript-to-cell assignment, rather than simple regeneration of the missing files. Consequently, submissions containing only transcripts and morphology could not be directly converted into the complete Xenium output structure required by our pipeline. This was a major contributor to the more than 200 samples that could not be automatically salvaged. Had the complete output generated by Xenium Onboard Analysis been deposited, these samples could have been processed directly without requiring dataset-specific reconstruction and reanalysis.

One of the limitations of the current pipeline is its inability to automatically process samples for which relevant data are uploaded directly to the GEO series page rather than provided through sample-level supplementary data records. These cases require manual downloading of the relevant files. In addition, when filenames do not provide a straightforward mapping between uploaded files and individual samples, manual supervision is required to establish the sample-level correspondence.

Importantly, most of the inconsistencies we encountered were not inherent to the Xenium data-generation or analysis process, but were introduced during data preparation and submission. Had the files generated directly by the Xenium software been deposited without renaming, restructuring, modification, or selective omission, many of the recovery and normalization procedures implemented in our pipeline would not have been necessary. The need for such a pipeline therefore arises primarily from the discrepancy between the standardized output produced by Xenium software and the heterogeneous form in which these data are deposited in GEO. This highlights the importance of preserving the original software-generated file structure when depositing spatial transcriptomics datasets in public repositories.

**Single-cell RNA reference atlas construction**

To construct reference atlases for cell type prediction, we collected publicly available human and mouse single-cell RNA-seq datasets with cell type annotations from the CellxGene Census (Megill *et al.* 2021). To minimize technical variation, only datasets generated using the 10x Genomics Chromium 3′ v3 platform were included. Protein-coding genes were retained based on GENCODE gene annotations.

Reference expression profiles were generated by aggregating single-cell expression data into pseudobulk profiles for each donor and annotated cell type using the R Seurat package (version 5.5.1)(Hao *et al.* 2021) . Specifically, gene counts were summed for each donor × cell type combination, normalized for library size, and log-transformed. Cell types represented by fewer than 1,000 cells were excluded. To reduce donor-specific batch effects, the normalized pseudobulk profiles were corrected using the removeBatchEffect function from the R limma package (version 3.6.9) (Ritchie *et al.* 2015), with donor as the batch variable while preserving biological differences between cell types. The corrected donor-level profiles were then averaged to generate a single reference expression profile for each cell type.

The resulting reference atlas contained 222 human cell types and 87 mouse cell types. Cell types were subsequently grouped into 18 human and 11 mouse broad cell-type categories based on hierarchical clustering and manual curation.

**Cell type prediction**

For each Xenium sample, the average gene expression profile of each cluster was calculated using the genes shared with the reference atlas. Spearman correlation coefficients were then calculated between each Xenium cluster and every reference cell type. Each cluster was assigned the reference cell type with the highest correlation together with its corresponding broad cell type.

Because Xenium datasets frequently contain relatively small targeted gene panels and are inherently noisy, anatomically implausible predictions occasionally occurred. To address this, we applied an anatomy-based filter that removed cell-type assignments incompatible with the tissue of origin while retaining biologically plausible cell types, including resident and infiltrating immune populations.

### SUPPLEMENTARY RESULTS

#### Database-wide quality trends

To explore trends in the quality of samples, we calculated several summary statistics for all 1,539 Xenium samples and used these as input to Principal Component Analysis (PCA):

- the number of cells (log10)
- the number of covered genes (log10)
- the average number of genes detected in each cell (log10)
- the average number of transcripts per cell (log10)
- the median number of genes detected in each cell (log10)
- the median number of transcripts per cell (log10)
- the total number of transcripts in the sample (log10)
- the fraction of non-zero values in the gene-by-cell matrix

PCA was conducted using the R function prcomp, including scaling and centering each feature to mean 0 and standard deviation 1. The resulting loadings of the 8 PCs are shown in Supplementary Table S1. In general, PC1 (which contained 59.7% of the variability in the data) had positive loadings for all quality indicators. In contrast, PC2 (22.0%) and other PCs were not consistently associated with all quality indicators. Figure S2 shows scatterplots of the 1,539 samples in the 2 first principal components, visualizing four representative quality indicators. As can be seen in the plots, PC1 is associated with the number of cells in each sample (Fig. S2A), the average number of transcripts per cell (Fig. S2B), and – to a lesser extend – the sparsity of the data (Fig. S2C). Other quality measures showed similar trends. The average number of genes detected per cell in a sample was positively correlated with PC1 and PC2 (Fig. S2D), with PC2 roughly separating samples by the number of genes included in the panels. Put together, we therefore concluded that PC1 is correlated with several properties that are generally understood to be signs of higher quality, and that it therefore can be used as a rough indicator of quality. Although this analysis is not meant to be interpreted as a rigorous quality assessment, it can be used to give users a rough indication of the quality of samples. Therefore, we decided to add this information to the sample table of the database.

### SUPPLEMENTARY FIGURES AND TABLES

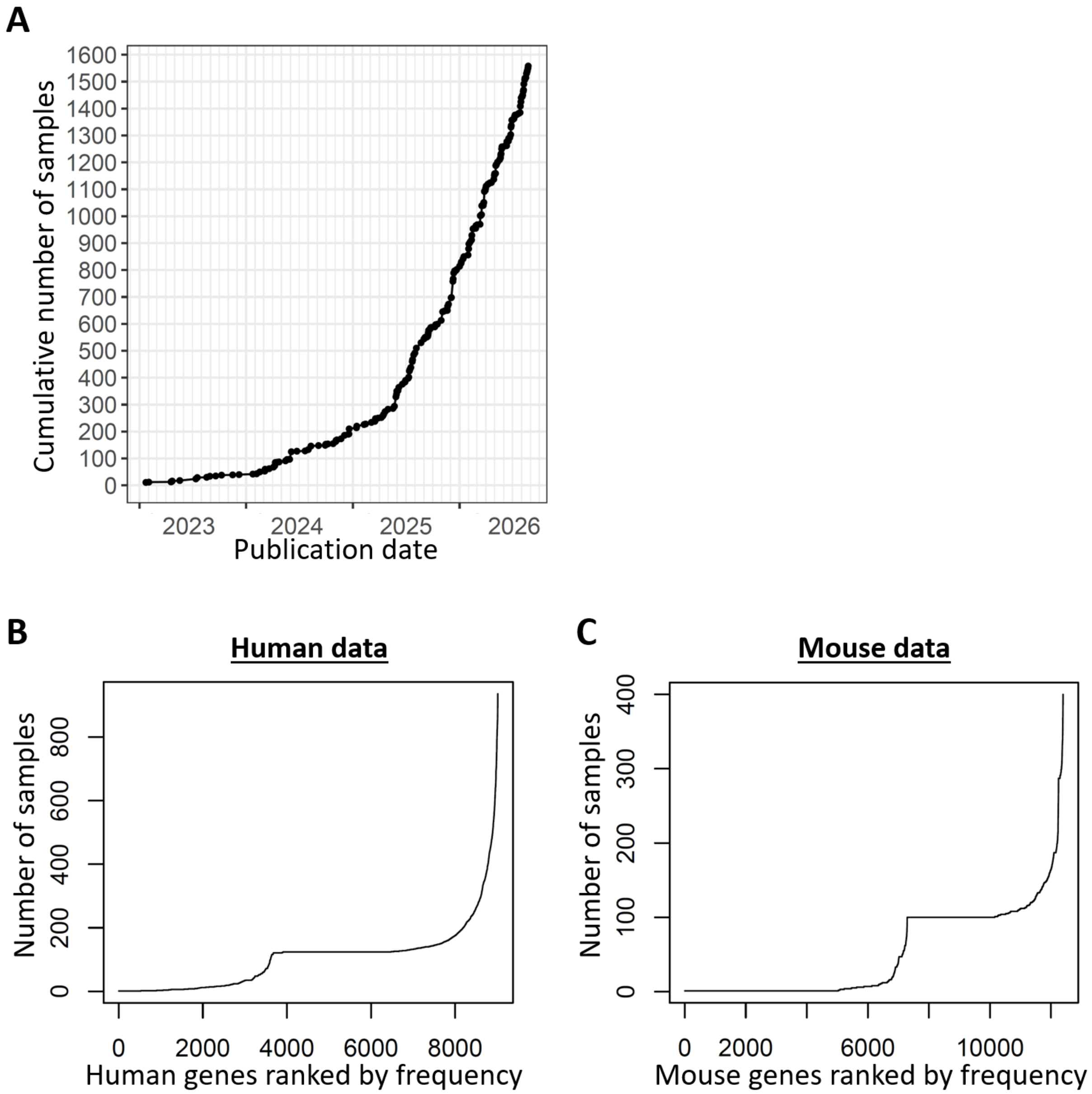


**Suppl. Figure S1.**

**(A)** The cumulative number of Xenium samples (as available in DeepSpaceDB) in function of their publication data.

**(B-C)** For human (B) and mouse (C) data, the number of samples in which each gene is included is shown in the y axis. The x axis shows the genes ranked by their frequency. In both cases, only a small number of genes is frequently included, and no gene is covered in all samples.

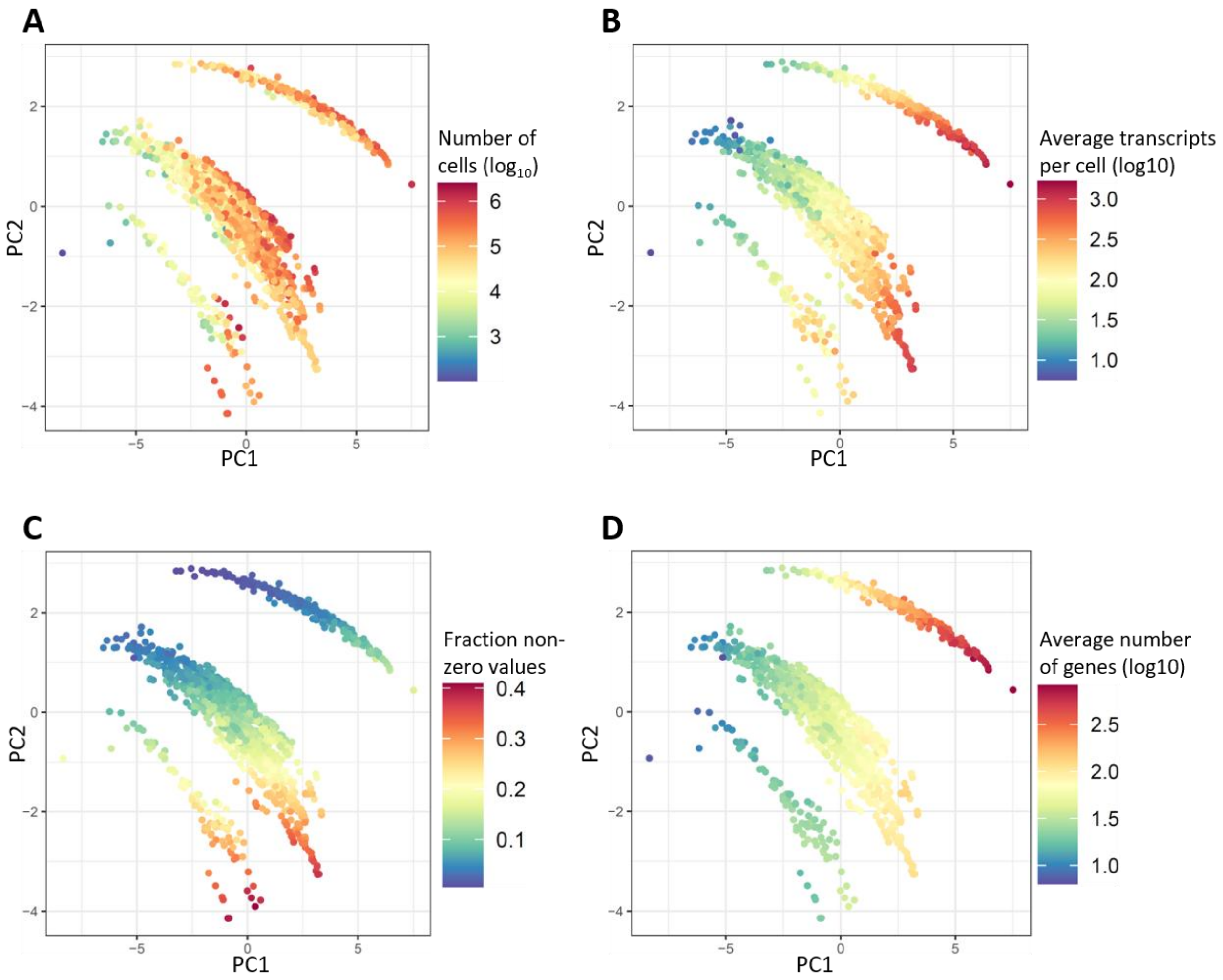


**Supplementary Figure S2.**

Scatterplots of PC1 (X axis) and PC2 (Y axis) show various quality indicators of the 1,539 Xenium samples.

**(A)** the number of cells ($log_{10}$ values).

**(B)** the average transcripts per cell ($log_{10}$ values).

**(C)** the fraction of non-zero values in the genes-versus-cell matrix. Lower values indicate higher sparsity.

**(D)** the average number of genes detected per cell ($log_{10}$ values)

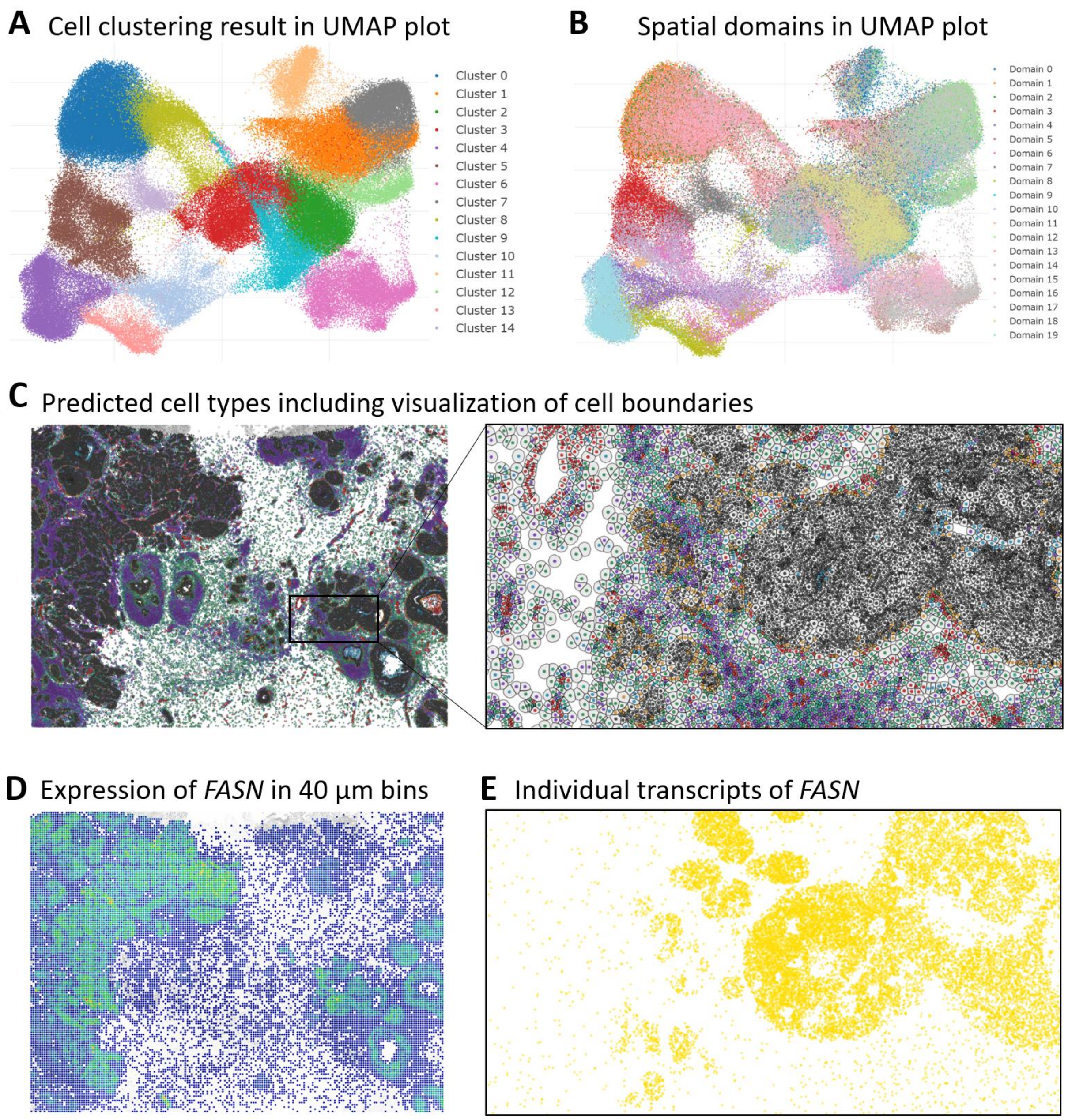


**Supplementary Figure S3.** This figure shows additional views of the same human breast cancer sample (sample ID DSIDX00001) shown in main Figure 3

**(A)** The cell clustering result shown in a 2D embedding.

**(B)** Spatial domains predicted by BANKSY shown in a 2D embedding.

**(C)** Cell boundaries are shown for a region of the tissue slice. Colors indicate predicted cell types.

**(D)** The expression pattern of *FASN* is shown in 40 µm bins. This is the same data as is shown on a single-cell resolution in Figure 3B in the main text.

**(E)** Individual transcripts for *FASN* are shown in yellow in the same region as is shown in panel **(C)**.

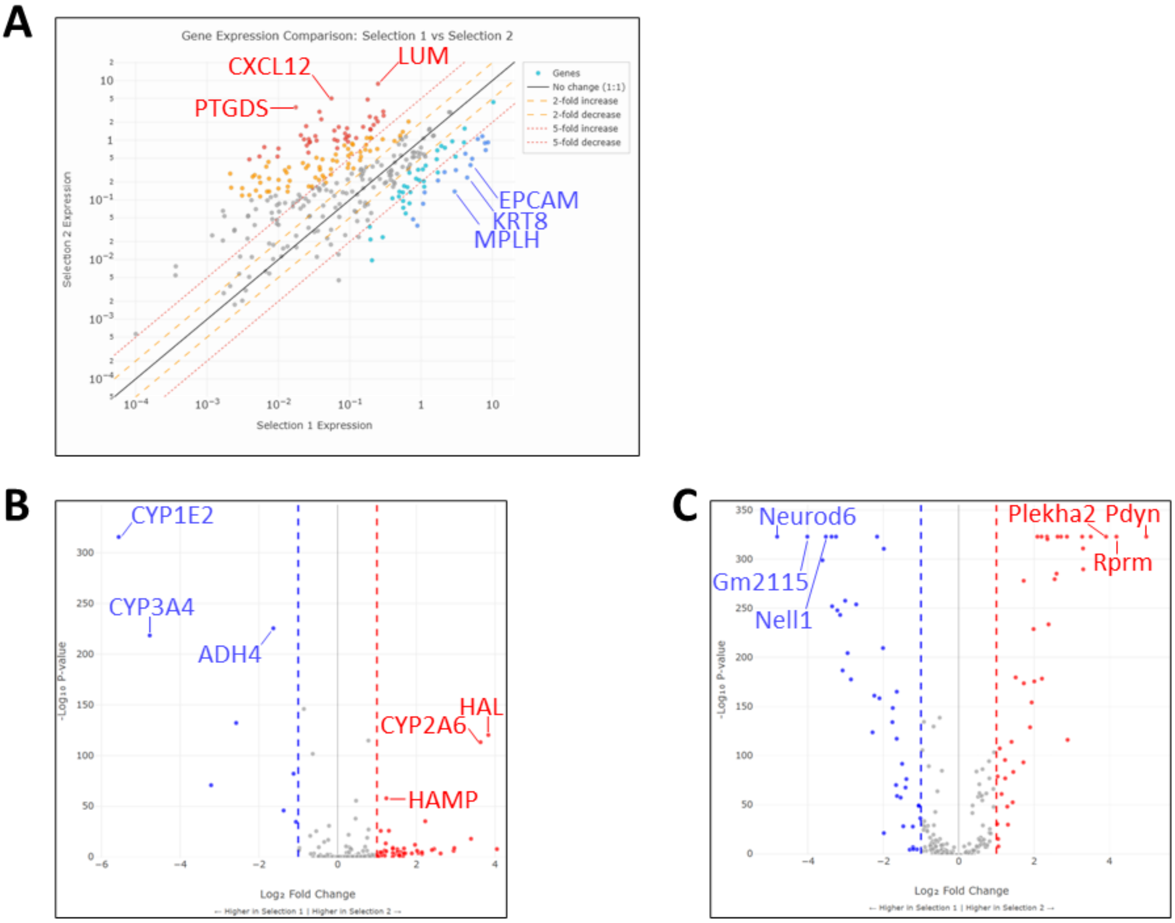


**Supplementary Figure S4.**

**(A)** Scatterplot of the average gene expression in the two selected regions of interest (ROIs) shown in Figure 5A. This is an alternative way of showing the same result as seen in the volcano plot shown in Figure 5B.

**(B)** A volcano plot of the gene expression levels in the 2 selected ROIs in Figure 5D. Genes with large differences in expression are indicated.

**(C)** A volcano plot of the gene expression levels in the 2 selected ROIs in Figure 5G. Genes with large differences in expression are indicated.

**Supplementary Table S1:** Loadings of the principal components. PC1 has positive loadings for all quality indicators.

| Feature | PC1 | PC2 | PC3 | PC4 | PC5 | PC6 | PC7 | PC8 |
|---|---|---|---|---|---|---|---|---|
| number of cells (log10) | 0.231 | -0.032 | 0.773 | -0.062 | -0.032 | 0.115 | -0.052 | 0.572 |
| number of covered genes (log10) | 0.265 | 0.599 | -0.059 | -0.118 | 0.709 | 0.207 | 0.091 | 0.000 |
| average number of genes detected in each cell (log10) | 0.438 | 0.114 | -0.150 | -0.382 | -0.452 | 0.119 | 0.639 | 0.000 |
| average number of transcripts per cell (log10) | 0.381 | -0.114 | 0.474 | 0.198 | 0.065 | -0.162 | 0.065 | -0.737 |
| median number of genes detected in each cell (log10) | 0.101 | -0.711 | -0.086 | -0.520 | 0.451 | 0.044 | 0.036 | 0.000 |
| median number of transcripts per cell (log10) | 0.231 | -0.032 | 0.773 | -0.062 | -0.032 | 0.115 | -0.052 | 0.572 |
| total number of transcripts in the sample (log10) | 0.265 | 0.599 | -0.059 | -0.118 | 0.709 | 0.207 | 0.091 | 0.000 |
| fraction of non-zero values in the gene-by-cell matrix | 0.413 | -0.182 | -0.258 | 0.504 | 0.185 | -0.515 | 0.216 | 0.360 |
